# Thermal variability interacts with infection load and the evolution of host-parasite defense

**DOI:** 10.64898/2026.07.20.739373

**Authors:** Rachael D. Kramp, Jennifer M. Cocciardi, Jason C. Walsman, Michel E. B. Ohmer

**Affiliations:** Department of Biological Sciences, University of Pittsburgh, Pittsburgh, PA 15260, USA; Department of Biological Sciences, University of Mississippi, University, MS 38677, USA; Earth Research Institute, University of California, Santa Barbara, Santa Barbara, CA 93106-9620; Division of Natural and Applied Sciences, Duke Kunshan University, Kunshan, Jiangsu Province, China 215316

## Abstract

Rising temperatures and unpredictable weather patterns are directly linked to emerging infectious diseases that threaten global biodiversity. Thermal variability affects host-parasite interactions, and understanding how animal hosts respond to temperature variability and parasite infection is paramount for conservation and for predicting zoonotic spillover. Hosts employ two strategies to defend against parasites: resistance (inhibiting or limiting infection) and tolerance (limiting the negative effects of infection). This observation raises a fundamental question: How does thermal variability affect the expression and evolution of resistance versus tolerance? Subsequently, how does the evolution of resistance versus tolerance influence parasite load dynamics? Here, we first review why temperature may differentially affect resistance and tolerance mechanisms, thereby altering host selection and eco-evolutionary disease outcomes. Second, to highlight the importance of these interactions, we present a model that illustrates key potential effects of temperature variability on host defense mechanisms. Our model demonstrates that temperature variability alone could drive lower infection prevalence and loads, but it also selects for host tolerance, ultimately leading to higher net prevalence and loads. We also find widely divergent outcomes depending on how temperature impacts defense strategies. These results highlight key areas for future empirical and theoretical work on the interactions among temperature variability, infection load, and host defense evolution.

## 1. Background

Climate change is increasing the frequency and intensity of temperature extremes and altering patterns of temperature variability (Q. Liu et al. 2026; Easterling et al. 2000; Yeh et al. 2009; Patel, Bonan, and Schneider 2024), as well as intensifying outbreaks of infectious diseases (T. E. Hector, Gehman, and King 2023). While the majority of research has focused on understanding the effects of a steadily warming climate, the physiological and immune responses of animals to temperature variation differ markedly from those to gradual temperature increases (Rebl et al. 2018; Vasseur et al. 2014; Terrell et al. 2013). Hence, predicting how animal hosts will respond to temperature variability, in addition to parasite infection, is paramount for conservation and for predicting zoonotic spillover (Ummenhofer and Meehl 2017; Gibb et al. 2020).

Changes in temperature variability can have strong physiological effects on ectothermic hosts because their body temperatures are regulated by environmental temperatures (Deutsch et al. 2008). Unlike gradual warming, increased temperature variability can more frequently expose individuals to conditions outside their optimal thermal environment, reducing performance and fitness (Vasseur et al. 2014; Buckley and Huey 2016). For example, increased temperature variation relative to a constant temperature or gradual warming can reduce the growth rates of ectotherms (Pisano, Kuparinen, and Hutchings 2019; Saxon, O’Brien, and Bridle 2018), and alter important thermal traits, including upper thermal tolerances and optimal temperatures (Clusella-Trullas, Blackburn, and Chown 2011; Paaijmans et al. 2013). This difference is important in the context of disease because host physiological performance is highly non-linear across temperature, meaning that exposure to temperature extremes and variations may disproportionately increase stress and reduce host defenses compared to gradual warming. While infection alone can alter host thermal physiology, for example, by reducing upper thermal tolerances (reviewed in (Tobias E. Hector, Sgrò, and Hall 2021) and increasing lower thermal tolerances (e.g., (Gehman, Hall, and Byers 2018; Linderman et al. 2012), increasing temperature variability may further compound physiological costs. Understanding how host populations will evolutionarily cope with the combined effects of temperature variability and disease, therefore, remains a key question. While animal hosts may have long generation times, their evolution can readily occur on time scales relevant to pressing issues of disease and climate change (e.g., Mountain Yellow-Legged frogs, *Rana sierrae,* take four years to reach reproductive age, but have evolved resistance to chytridiomycosis in a few decades (Byrne et al. 2026)).

All host defenses against parasites (whether via behavioral or physiological mechanisms) can be divided into two broad strategies: resistance and tolerance (Roy and Kirchner 2000; Råberg, Graham, and Read 2009). These broad categories classify defense strategies according to how the host’s defense affects the parasite’s fitness. Resistant hosts improve their fitness at the expense of parasite fitness, such as by preventing infection, recovering from infection more quickly, or limiting infection load (the number of parasites per host). In contrast, tolerant hosts improve their fitness without harming parasite fitness, e.g., by surviving significant parasite burdens without dying, which can even increase parasite fitness (Best, White, and Boots 2008; M. R. Miller, White, and Boots 2005). When considering parasites that replicate directly within a single host, resistance reduces infection load, thereby increasing host survival and recovery rates. At the same time, tolerance allows hosts to survive higher loads. The course of infection within a single individual directly informs population-level infection dynamics and ultimately the selective pressure within a given population (Handel A. 2015; Mideo, Alizon, and Day 2008). When considering the number of parasites (infection load) infecting a host, there is a positive trend: as the infection load increases, transmission also increases. Therefore, resistance traits reduce parasite prevalence by directly reducing parasite growth and ultimately reducing parasite fitness. In contrast, tolerance only mitigates parasite damage, potentially allowing extended and more intense infectious periods and thereby increasing, rather than decreasing, parasite prevalence within the population (Best, White, and Boots 2008; Boots and Bowers 1999). The evolution of resistance versus tolerance strategies can reshape host-pathogen interactions, alter transmission dynamics, and shape selection on parasites. Understanding how temperature affects these defense strategies is, therefore, a priority when considering how a changing climate affects infectious disease (Altizer et al. 2013). Yet most research focuses on temperature-disease effects at either the individual-host or population level, rarely both (Kirk, O’Connor, and Mordecai 2022). To address this gap, we consider the severity of temperature variability and disease impacts on the host relative to those on the population, using the key mechanism of infection load (Wilber et al. 2022).

Although resistance and tolerance both commonly evolve to combat a variety of infections in natural systems (Råberg 2014; Råberg, Sim, and Read 2007; Gehman, Hall, and Byers 2018; Holand et al. 2019), they are generally thought to trade off against each other (Råberg, Sim, and Read 2007; Best, White, and Boots 2008). The trade-off has been empirically demonstrated across multiple species (Klemme and Karvonen 2017; Kramp et al. 2026). In fact, mathematical modeling shows that greater environmental variability, specifically through a periodic host birth rate, can increase infected population density, increasing the force of infection and selecting for greater investment in host tolerance. Investment in resistance that prevents infection may respond to increased force of infection by increasing or decreasing as environmental variability increases (Ferris and Best 2019); such resistance is known to be selected for or against when the force of infection rises, depending on conditions (Jason C. Walsman et al. 2023). This model indicates that the two strategies respond in qualitatively opposite ways to the same environmental pressures, a divergence that holds across both constant and variable environments. However, concurrent evolution of resistance and tolerance has been theorized and demonstrated in plants and birds (Restif and Koella 2004; Singh and Best 2021; Bonneaud et al. 2019). Furthermore, there is evidence that tolerance can evolve without a reduction in antibody-specific antigen levels, thereby maintaining resistance while tolerance evolves (Hayward et al. 2014). Therefore, as climate change continues, host evolution may be driven by both parasite infection and temperature fluctuations, leading to increases or decreases in parasite prevalence within populations.

Evolutionary trajectories of resistance and tolerance are shaped by ecological context and pathogen pressure. When a novel pathogen enters a naive population, and the transmission risk and infection cost are high, resistance is likely to provide optimal protection while preserving host fitness. This strategy reduces the risk of infection and maintains reproductive potential by limiting infection risk and maintaining mate attraction. Behavioral immunity, a type of avoidance response, can rapidly shift transmission dynamics (Shakhar 2019; Poulton and Ellner 2025), and individuals who can avoid or suppress infection have greater mating success. However, when parasite prevalence is high and infection becomes nearly unavoidable, resistance may no longer be the optimal strategy (Jason C. Walsman et al. 2023). In these scenarios, tolerance is more advantageous, as it reduces the fitness costs of inevitable infection (Budischak and Cressler 2018). Importantly, tolerance differs fundamentally from resistance in its environmental feedback: both strategies incur energetic and physiological costs that shape their evolutionary trajectories; resistance entails immune activation and associated pathology (Knutie et al. 2017; Amo et al. 2021), manifesting as reduced growth or reproductive output; tolerance can mean enduring chronic infections and, in some cases, reduced lifespan (Ayres and Schneider 2012). Life-history traits further mediate these strategies: species with rapid reproduction may favor resistance, while longer-lived species may benefit more from tolerance or a combination of both (Sears, Snyder, and Rohr 2015).

Previous work has demonstrated that warming can increase parasite prevalence, and that environmental factors, such as temperature, resource availability, and moisture, play a crucial role in modulating parasite transmission and host defense investment (Rosa et al. 2022; McNew et al. 2019). Using a high-resolution 9,600-year record of trematode parasite traces in bivalve hosts from the Holocene Pearl River Delta, Huntley et al. (2014) found that parasite prevalence increases during periods of warming, and cannot be explained by host abundance or salinity changes. The authors therefore predict that warming and climate-related alterations will result in patterns similar to those observed in the past (Huntley et al. 2014). The exact reason for the increase in parasite prevalence is unknown. Higher temperatures may have increased parasite reproduction, resulting in greater damage to the host during warming events, leading to increased mortality and potentially reduced transmission rates. Under these circumstances, if tolerance enhances hosts’ ability to mitigate parasite damage, then hosts with high tolerance should experience positive selection. Ultimately, a population with more tolerant individuals should also have a higher prevalence, even without an increase in host abundance. Tolerance provides a possible explanation for the increased prevalence observed in fossils during a warming climate. Environmental pressures imposed by climate change could differentially impact resistance and tolerance and, therefore, parasite prevalence, thereby directly contributing to spillover risk (M. R. Miller, White, and Boots 2005; Martin R. Miller, White, and Boots 2006; Guth et al. 2022). Despite these major implications, it remains unclear whether hosts allocate resources differently across alternative defense strategies in response to increased temperature variability.

Theoretical and empirical research has shown that fluctuating environments drive changes in host-parasite evolution (Ferris and Best 2019; Duncan et al. 2017; Harrison et al. 2013; Donnelly et al. 2013; Blanford et al. 2003; Koelle, Pascual, and Yunus 2005; Sorrell et al. 2009; Ferris and Best 2018), most of which focus solely on parasite evolution. Among the studies above that do examine host evolution, most focus exclusively on host resistance and the immune system. Here, we present a conceptual review examining the intersection of temperature variability, host defense—particularly the host immune system, which is key to the evolution of host resistance—and parasite load, an area that has yet to be thoroughly investigated (**Concept 1, Dark blue center**). Furthermore, we used a mathematical model based on data from a focal host-parasite system, the Mountain Yellow-legged frog (*R. sierrrae*) and the amphibian chytrid fungus, *Batrachochytrium dendrobatidis (Bd)*, to explore how daily temperature variability shapes the evolutionary trajectories of the two host defenses. By tracking interacting variables for the densities of susceptible and infected hosts and their loads, as well as environmental parasites (zoospores), we identify important possible outcomes in disease dynamics, such as parasite prevalence and population dynamics, over discrete daily time steps.

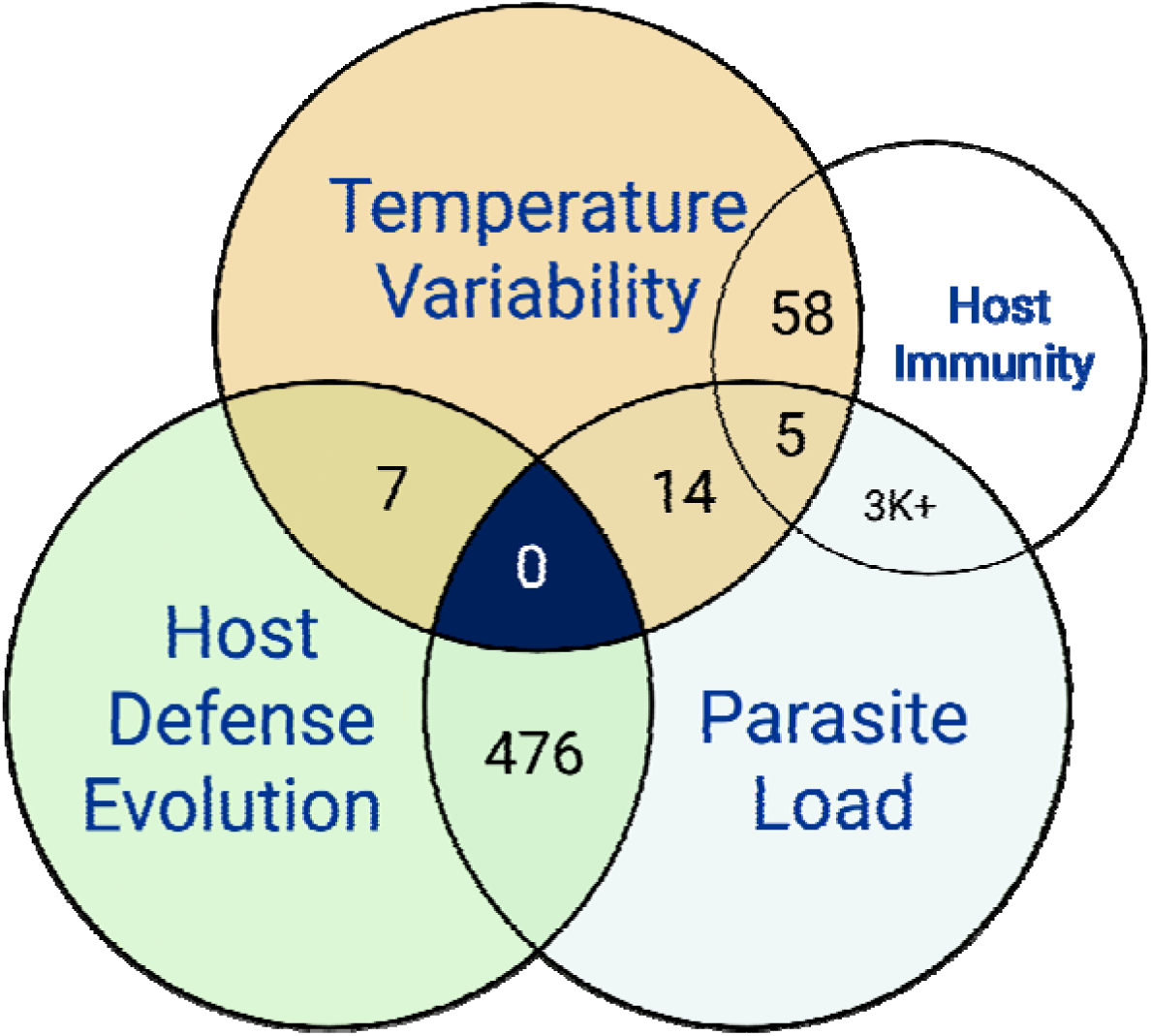

**Concept 1. Mapping the research gap in the evolution of host defense during increased temperature variability and parasite load or prevalence.** The Venn diagram illustrates the intersection of three key domains: Host defense strategies (Resistance or Tolerance), Temperature Variability, and parasite load or prevalence. While previous studies have examined how temperature variability affects the host immune system, to our knowledge, there is no research on the evolution of host defense under the dual stress of specifically temperature variability and parasite infection. However, it has been suggested within the literature (Thomas R. Raffel et al. 2015). These broad classes of host defense strategies provide a framework for predicting parasite prevalence within populations, which informs host conservation strategies and potential zoonotic spillover. Within our keywords, we did not include the general term “environmental variation,” which has a larger body of literature and includes biotic and abiotic factors. The supplementary information contains keywords and searches performed on Web of Science. Created in BioRender. Kramp, R. (2026) https://BioRender.com/i3cosdg.

## 2. Impacts of temperature extremes and infection on host physiology and defense strategies

Extreme temperatures have been identified as the most significant selective force on species in the context of ongoing global climate change (Gutschick and BassiriRad 2003; Grant et al. 2017). Animals are being increasingly exposed to temperatures outside their thermal optima (Kirk, O’Connor, and Mordecai 2022). Several parasite species have been shown to reduce the host’s ability to endure extreme temperatures; however, the mechanisms of action remain unknown (T. E. Hector, Gehman, and King 2023). Parasite infection can decrease a host’s thermal ranges and optima (Gehman, Hall, and Byers 2018). While not inherently parasitic, a host’s microbial communities can also alter the host’s ability to withstand heat stress and profoundly change the host’s physiology in response to it (Fontaine and Kohl 2023). This could be one way in which some parasites disrupt a host’s thermal tolerance (J. D. Li et al. 2024). Furthermore, even when hosts carry similar loads, warmer climates appear to result in greater host damage, effectively increasing parasite virulence (Debes, Gross, and Vasemägi 2017). These infection-related impacts are a critical way in which temperature interacts with host defenses, as resistant hosts can recover from infection more quickly, thereby selecting for stronger resistance when infection synergizes with temperature to harm hosts. This prediction depends on recovered hosts having resilient thermal ranges that return to pre-infection levels or remain narrower after infection. For example, one such study found that tropical lizards exhibited increased thermal tolerance following antiparasitic treatment (Bakewell et al. 2025). However, to our knowledge, the resilience of thermal ranges remains underinvestigated in many host-parasite systems.

Defense traits, such as resistance and tolerance, interact with many physiological mechanisms and may influence thermal range tolerance. This may indicate that tolerant traits confer an advantage when facing stress that requires similar cellular repair. Heat stress could potentially amplify the costs of resistance by increasing damaging inflammation (Heled, Fleischmann, and Epstein 2013), 2013; (Quance and Travisano 2009)) and can suppress aspects of host immunity, thereby increasing host susceptibility to parasites (Zheng et al., 2016), and possibly making resistance traits less effective (Meng et al. 2013). Conversely, tolerant hosts expend less energy on immune responses and may be better equipped to maintain their physiological stability under heat stress. Known host tolerance mechanisms, such as preventing or reducing immunopathology or altering thermoregulation (e.g., by shifting the host’s thermostatic set point) (McCarville and Ayres 2018; Schieber and Ayres 2016), could be critical for tissue protection during heat-induced hyperstimulation of proinflammatory cytokines (Schieber and Ayres 2016; E. Liu et al. 2012). Likewise, tolerant traits could be more labile in response to environmental changes than resistant traits, a key advantage when facing multiple stressors (McNew et al. 2019). Thus, host genotypes that are more tolerant of parasites may also be better able to survive and function under unusually high temperatures.

However, if animals under heat stress reduce their resource intake (due to physiological impairment or behavioral change), selection may favor resistance strategies. Heat stress has complex effects on animal physiology; evidence indicates that it can impair gut permeability and induce anorexia (Cao et al. 2021; Kanjanapruthipong, Junlapho, and Karnjanasirm 2015). Decreased resource acquisition can reduce tolerance effectiveness, and some evidence suggests that low resource availability increases antibody-mediated immune responses in frogs, whereas high resource availability appears necessary for frogs to recover energy lost to parasite infection and increase tolerance (Knutie et al. 2017). Thus, if animals reduce their resource intake under heat stress, resistance traits could increase within the population. Development, in combination with limited resource availability, can further mediate this balance, as investment in growth or reproduction may limit the energy available for costly immune responses or tissue repair (Doeschl-Wilson et al. 2009; Råberg 2014).

## 3. How does environmental change impact infection loads within hosts?

Thermal variability often benefits the pathogen over the host, as pathogens are typically much smaller and can acclimate more quickly to changing conditions (Rohr and Cohen 2020; Cohen et al. 2017). As a result, thermal variability can increase infection loads, particularly if temperature changes shift hosts away from their thermal optimum. Indeed, increased thermal variability increased infection prevalence and burden in a *Drosophila*-microparasite system, which was opposite to the predictions of a mechanistic disease transmission model that incorporated acclimation responses of hosts and pathogens (Krichel et al. 2023). The authors hypothesized that this was the result of disease processes interacting across scales (from the individual to the population), which were not captured by their transmission models incorporating metabolic scaling theory. Furthermore, thermal variability can prompt the host to prioritize upregulating metabolic processes over immune function (Jörn P. Scharsack et al. 2021), thereby increasing infection burdens. In addition, previous acclimation to warmer temperatures results in higher infection loads following a decrease in temperature in the amphibian-chytridiomycosis system (Thomas R. Raffel et al. 2015; Noelker et al. 2024).

As extreme events become more prevalent, we often see increases in infection prevalence and intensity following heat waves and cold snaps. In the *Daphnia magna–Ordospora colligata* host-parasite system, experimental heat waves increased infection burdens by up to 13-fold over baseline, but this depended on the timing, duration, and magnitude of the heat wave, as well as the starting temperature (McCartan et al. 2025). Cold-snaps, on the other hand, resulted in either increased or decreased infection burden, depending on the baseline temperature, with greater increases in load following a cold-snap at the highest baseline temperature (McCartan et al. 2024). This highlights the importance of thermal variability, rather than just thermal extremes, in driving infection dynamics. These impacts can be partially mediated by acclimation, as previous work has found that acclimation to warmer temperatures can decrease pathogen burdens and increase survival (Scanes et al. 2023).

## 4. How does environmental change impact the immune system of ectothermic hosts?

Current research investigating the effects of temperature variability on the host immune system has shown that rapid temperature changes are generally associated with immune suppression. In red-spotted newts, both short-term temperature lags and seasonal acclimation impact immune parameters in field-collected animals (T. R. Raffel et al. 2006). In particular, lymphocyte levels were reduced after rapid warming in the spring, supporting a lag effect, while neutrophil, eosinophil, and lymphocyte numbers decreased in the fall, supporting a seasonal acclimation effect. Furthermore, Terrell et al. (2021) found that hellbenders are susceptible to reduced growth and disrupted immune function at both warm (>22°C) and cool (<14°C) temperatures, but that bacterial-killing ability is reduced during temperature changes (Terrell et al. 2021). Bozinovic et al. (2013) found that the immune system was most affected by extreme shifts in temperature variability in beetles, with greater variation in the host’s ability to kill bacteria (Bozinovic, Catalán, and Kalergis 2013), showing that both short-term and seasonal temperature shifts can affect both innate and adaptive immune cells. Similarly, teleost fish immune function is greatly influenced by temperature and is optimal at intermediate temperatures within their range (Jörn Peter Scharsack and Franke 2022). Adaptive and specific immune responses appear to be suppressed by cold temperatures, forcing teleost fish to rely on innate immune responses (Bly and Clem 1991; Rijkers et al. 1981; Kyprianou et al. 2010). Amphibians exposed to experimentally variable temperatures were more susceptible to chytridiomycosis, a fungal infection caused by the pathogen *Batrachochytrium dendrobatidis (Bd;* (Greenspan, Bower, Webb, et al. 2017; Thomas R. Raffel et al. 2015, 2013). The physiological shift was more taxing for animals that were acclimated to warm temperatures. This finding aligns with the observation that amphibians are more susceptible to *Bd* infection following decreases in temperature than increases and underscores the consensus that increased temperature variability associated with climate change may increase the impact of infectious diseases. A shift from 20°C to 8°C in common carp (*Cyprinus carpio*) induced a delay in the primary antibody response, and the secondary immune response was present from 24°C to 20°C, but was lost at 18°C and below (Rijkers et al. 1981; Rijkers, Frederix-Wolters, and van Muiswinkel 1980). These studies indicate that temperature variability will have an important impact on the host’s susceptibility to pathogens and, likely, on the spread within populations.

Over time, we should see hosts evolve in response to increased temperature fluctuations due to climate change. These trends may indicate that host resistance (immune system) is becoming less effective due to increased temperature variability, thereby favoring more tolerant individuals. However, this is a complex interaction, and some research suggests that it may have opposite effects. Terrell et. al. (2013) found that temperature variation increased the immune system’s ability to kill bacteria in salamanders. Specifically, they surmised that an elevated innate immune defense might be adaptive in an unpredictable environment (Terrell et al. 2013). Overall, warm temperatures often increase immune responses, while cool temperatures suppress them, with many vertebrates showing thermal dependence of constitutive defenses (Butler et al. 2013), and upregulation of immune genes with warm acclimation (Jiaying Li et al. 2025). However, at thermal extremes, some species are immunocompromised (Jörn Peter Scharsack and Franke 2022).

## 5. Case study: Impacts of temperature variability on Bd infection load and the evolution of defense strategies in Mountain yellow-legged frogs (*Rana sierrae*)

In this paper, we use the amphibian-Bd host-pathogen system as a case study to explore the impacts of daily temperature variability on infection load and the corresponding evolution of resistance and tolerance defense strategies. In particular, we parameterize our model using data from mountain yellow-legged frogs (*R. sierrae,* hereafter MYL frogs) in the Sierra Nevada of California. Populations of this species have been monitored for Bd over the past decade, documenting the species’ decline and subsequent recovery in parts of its range (Knapp et al. 2016). Based on an ongoing translocation and reintroduction program, there is evidence that some populations are persisting with Bd, and when those populations served as source populations in translocations, translocation success was higher (Knapp et al. 2024), as well as genetic evidence for evolution driving recovery of some populations (Byrne et al. 2026). This provides evidence of the evolution of resistance and/or tolerance to Bd in these populations. Importantly, published experimental work provides temperature-specific parameters, such as survival and on-host Bd growth, as a starting point for modeling (Wilber et al. 2016).

### Model methods

Pathogen load mechanistically links infection severity and heterogeneous infection outcomes for individual hosts to population-level dynamics of disease spread and the suppression of host abundance. Following (Jason Cosens Walsman et al. 2026) we model the density of susceptible hosts, the density of infected hosts with a log pathogen load, i.e., the natural log of the number of pathogens on an infected host, in some range *x* to *x*+*dx*, and the density of environmental pathogens at each discrete time step *t*. New susceptible hosts are produced by infected and susceptible hosts at a per-capita rate that declines with host density. Susceptible hosts may become infected at a rate that depends on the density of environmental pathogens, and newly infected hosts have load *x* drawn from a normal distribution with some mean and standard deviation. Pathogen load grows with some variability, so that some proportion of infected hosts experience decreasing loads, but tend to grow over time and approach a within-host carrying capacity. Note that the distribution of how many hosts have a certain log pathogen load, while following a normal distribution for new infections, changes dynamically and is not constrained to follow a normal distribution. Hosts with a higher pathogen load at time *t* shed more pathogens into the environment, have a higher probability of death before the next time step, and have a lower probability of recovering back into the susceptible class at time *t*+1 (see conceptual diagram in Figure 1).

**Figure 1.**
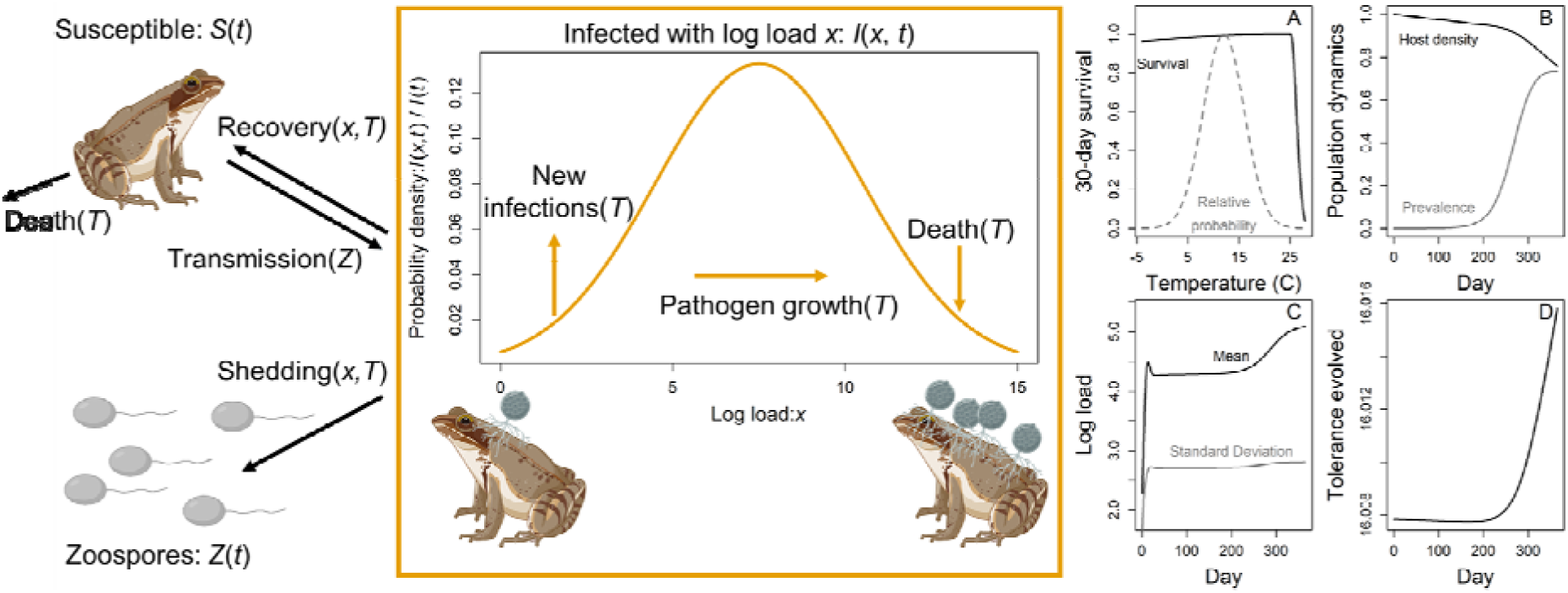
Temperature variability impacts survival and epidemic dynamics. The conceptual diagram illustrates the state variables of susceptible host density, *S*(*t*), the density of hosts infected with some load in the range *x* to *x*+*dx*, *I*(*x*,*t*), and the density of zoospores in the environment *Z*(*t*). Arrows show rates and the dependencies they can have on load, *x*, zoospore density, *Z*, and temperature, *T*. Gold arrows show how processes tend to alter the distribution of load as new infections primarily contribute low load infections, pathogen growth moves infections from lower loads to higher loads, and infected host death primarily reduces the number of high load infections. (A) Host survival probability (black curve) declines steeply at high temperatures. Here we show the survival probability for a susceptible host for one day raised to the 30th power, i.e., the probability of a susceptible host surviving thirty days at a given, constant temperature. The gray curve shows the relative probability density for a day having a given temperature for a mean temperature of 12 °C and a standard deviation of 4 °C. (B) Over the course of an epidemic, host population densities (black) decline as infection prevalence (gray) rises. Temperature fluctuations that affect mortality have minimal impacts on these dynamics. (C) The mean (black) and standard deviation (gray) of log load increase to stable levels during the epidemic (D) while hosts display very little evolutionary change in tolerance.

We model host evolution as competition among clonal host genotypes for simplicity. Clonal evolution is a common modeling method to isolate the effects of differential fitness among genotypes; including details of sexual reproduction, etc. would likely have a small impact on the results (see a discussion of this the context of amphibian tolerance evolution to amphibian chytrid; (Streipert et al. 2025)) but would also significantly complicate the model while requiring many additional assumptions (e.g., about inheritance and dominance patterns). We do allow mutation among clones as susceptible and infected individuals reproduce with a small chance of producing any other genotype. Genotypes vary in their defense, exhibiting either 1) the wild-type traits, 2) a lower pathogen growth rate than the wild-type (resistance), or 3) better survival while infected than the wild-type (tolerance). Genotypes with stronger defense of either form have lower fecundity, and the curvature of this defense-fecundity tradeoff selects for intermediate defense or, in the absence of disease, the wild-type. We simulated to find the endemic equilibrium of the wild-type in the presence of disease. Evolutionary simulations are then initialized at the susceptible density, infected density, load distribution, and zoospore density corresponding to that equilibrium with no mutation and no temperature variation; the initial host population is then set to 91% wild-type and 1% of nine other genotypes, all with the same distribution of load at *t* = 0 as found for the wild-type equilibrium and then the evolutionary simulation is run with mutation, evolution, possibly temperature variation, etc.. We also conducted simulations with the wild-type genotype starting at its no temperature variability, endemic equilibrium simulated with temperature variability to disentangle the effects of temperature variability alone from those of temperature variability and host evolution (Figs. S1 and S2). To distinguish the effects of temperature variability from high temperatures, we also simulated rising mean temperatures with no variability (see Figs. S3 and S4) across a range of temperatures chosen to impose the same mortality on an uninfected, wild-type host genotype at the highest mean temperature with no variability as suffered at the highest variability with no increase in mean. The endpoint of simulations were approximated as averages over the final 365 days of a 4 x 10^4^ day (∼110 years) simulation to allow time for host evolutionary and ecological dynamics to approach their endpoint, a process slowed by temperature stochasticity and the relatively slow generations of the focal hosts for whom we chose parameter values, Mountain Yellow-Legged Frogs (*Rana sierrae*).

We incorporate daily temperature fluctuations to determine the impact of temperature variability on disease spread and host evolution. Mean temperature stays the same, but the temperature on day *t* is drawn from a normal distribution with some fixed mean and standard deviation (Fig. 1A gray curve). For simplicity, temperature is drawn independently from this distribution for each day (but see Discussion regarding autocorrelation in future work) and influences the rates that determine the densities of susceptible hosts, infected hosts with each load value, and zoospores on the next day. In all model cases, we consider host survival probability to follow a thermal performance curve (TPC, black curve in Fig. 1A), particularly so that high temperatures are dangerous for host survival; for infected hosts, this survival probability curve as a function of temperature is multiplied by survival probability as a function of pathogen load, further reducing survival probability. In some model cases, we also consider temperature dependence in the initial load of newly infected hosts, pathogen growth rate, pathogen shedding rates, and recovery rates (see rates that can depend on *T* in conceptual diagram part of Fig. 1). Additionally, host defense and infection may interact with the survival TPC; in one case, more tolerant hosts have a higher heat tolerance, in another case infection reduces heat tolerance, and in a third case both effects are active. We determine how temperature variability impacts the spread of disease, suppression of host populations, and evolution of host defenses (See Fig. 1B-D).

When possible, we parameterize our model with biologically relevant values from the focal system of Mountain Yellow-Legged Frogs and a deadly fungal pathogen (*B. dendrobatidis*). Empirical measurements strongly inform fecundity, survival without disease, transmission rate, initial loads with their positive temperature dependency, within-host pathogen growth rate with its positive temperature dependency, recovery rate with its negative temperature dependency, shedding rate with its positive temperature dependency, and the probability of survival of environmental pathogens (see Appendix for values and citations). The TPC of survival is approximated based on a critical thermal maximum of 34 □ found for a related species, *Rana cascade* (Pottier et al. 2025), and daily survival probability of 0.9898 found for *Rana mucosa* at 26 □ (Cheryl Briggs, personal communication; see Fig. 1A for resulting TPC). The fecundity costs of defense are set so that hosts evolve an intermediate degree of defense, and that degree of defense is sensitive to model parameters. The effects of infection on the survival TPC and tolerance on the survival TPC are illustrative values chosen to qualitatively demonstrate different possible outcomes and to show that, in the scenario when both tolerance and infection affect the TPC, the most tolerant genotype has a slightly higher heat tolerance when infected than a wild-type uninfected host.

### Model results

Temperature variability can have conflicting effects on epidemic outcomes, depending strongly on whether and how hosts evolve. Temperature variability does not greatly affect host survival or the course of epidemics and host evolution (orange curve in Fig. 2). However, when host defense interacts with host’s heat tolerance, then temperature variability can greatly alter outcomes for host evolution, epidemics, and host population density. When tolerance only increases heat tolerance, little changes as hosts experience few deaths from heat (blue curves in Fig. 2B-F is similar to orange curves in Figs. 2B-F). When infection lowers heat tolerance, more hosts die, lowering host density (Fig. 2B green curve) and prevalence (Fig. 2C green curve) and weakening selection for host defense (Fig. 2E green curve). But when infection lowers heat tolerance *and* pathogen tolerance correlates with higher heat tolerance, temperature variability has large negative impacts on host density (Fig. 2B yellow curve) and large positive impacts on prevalence (Fig. 2C yellow curve). This arises because hosts evolve a high degree of pathogen tolerance to help with heat tolerance (Fig. 2E yellow curve), driving an increase in pathogen prevalence and loads (Figs. 2C-E), leading to high shedding and disease spread. The evolution of tolerance causes many hosts to suffer infection and infection-reduced heat tolerance, explaining the strong reductions in host population density.

**Figure 2.**
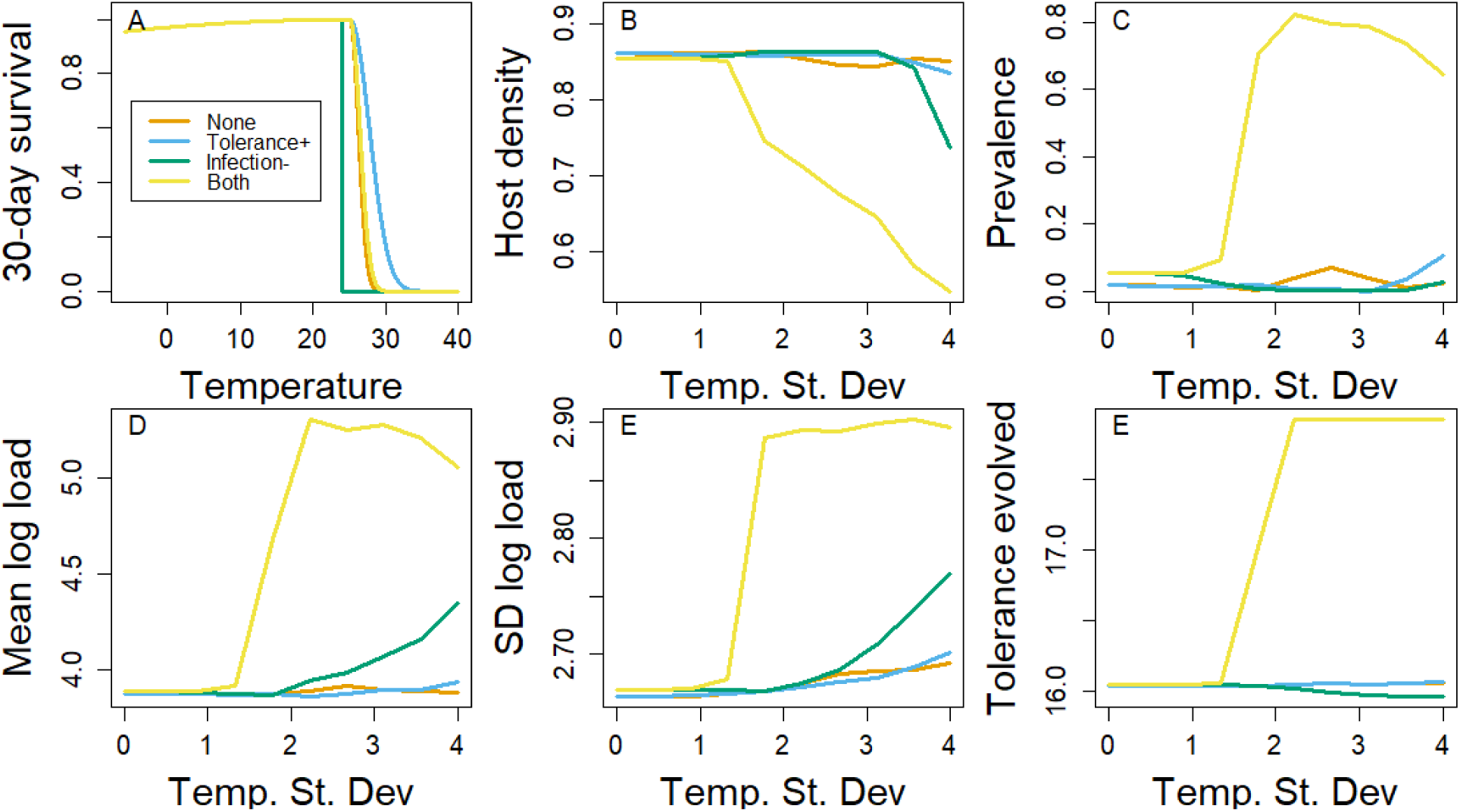
Thermal performance-disease interactions alter how temperature variability impacts host defense evolution. (A) We show a default thermal performance curve for the host probability of surviving for 30 days (orange) as well as the scenarios in which more pathogen-tolerant genotypes have higher heat tolerance (blue), infected individuals have lower heat tolerance (green) or both (yellow). For each scenario, the curve shows the TPC of the most tolerant genotype while infected (though in some scenarios, e.g., orange, genotype and infection do not impact this TPC). We then show the impact of the standard deviation of daily temperature (x-axis) on the interacting, long-term outcomes of (B) host density, (C) infection prevalence, (D) mean log load, (E) the standard deviation of log load, and (F) tolerance evolved

## 7. Discussion

The evolutionary response of a host to the dual stressors of infectious disease and temperature variation has yet to be tested. Predicting how animal hosts will respond to temperature variability and parasite infection is paramount for conservation and predicting zoonotic spillover (Ummenhofer and Meehl 2017; Gibb et al. 2020). We reveal fundamental gaps in research on these topics: intersections between defense evolution, temperature variability, and infection load. There are likely interactions between host defense against parasites and temperature that are well supported empirically: infection can shift thermal performance curves, and high temperatures exacerbate resistance costs (Gehman, Hall, and Byers 2018; Greenspan, Bower, Roznik, et al. 2017; T. E. Hector, Gehman, and King 2023; Will et al. 2025). We need to empirically test these relationships to predict host evolution, as our model findings emphasize. Under certain conditions, tolerance can evolve within the population, ultimately increasing pathogen prevalence. When thermal performance interacts with infection/defense, outcomes are dependent on the impact of infection on thermal tolerance; if infection narrows the host thermal performance curve (TPC), resistance may be favored in some cases or disease spread may be curtailed enough to weaken selection for host defense (as we found). But if infection broadens or aids with maintenance of the TPC, tolerance is favored. If clearing the infection also results in hosts regaining their normal TPC, this would favor hosts that could clear the infection under temperature variability, selecting for resistance. However, if tolerant hosts have a greater thermal breadth (e.g. Porras et al. 2020), thermal variability in the presence of pathogens can select for tolerance.

The hypothesis that tolerance broadens the TPC is untested. Host thermoregulation has been proposed as a tolerance-based trait, and evidence suggests that fever and heat shock induce cytoprotective and tissue-protective responses that help limit tissue susceptibility to damage (Schieber and Ayres 2016). Hector et al. found evidence of a marginally weak negative association between infection burden and temperature tolerance (T. E. Hector, Gehman, and King 2023). The authors note that further research is needed. However, this could be because tolerance broadens the temperature tolerance range during infection, i.e., more tolerant hosts survive higher temperatures and endure higher parasite loads. Therefore, eliminating the strong correlation between parasite infection and the host’s decreased ability to tolerate heat. The thermal breadth of tolerant vs. resistant hosts has not been empirically tested. Here, we demonstrate that if this is a tolerance trait, increased temperature variability due to climate change may increase the frequency of tolerance within populations. This could lead to higher parasite burdens and prevalence, thereby increasing the risk of spillover (Guth et al. 2022; Seal, Dharmarajan, and Khan 2021). Tolerant hosts are also associated with more virulent pathogen spillover; therefore, this could have major implications for public health and livestock (Brook et al. 2023; Guth et al. 2022). On the other hand, resistant individuals may have lower thermal tolerance. Some research suggests that artificially selected resistant individuals have lower survival at both low and high temperatures compared to controls in flies (Luong and Polak 2007).

In the context of a more variable climate, it is important to consider that resistant and tolerant traits are most likely plastic and depend on their environment (Sternberg et al. 2012). Phenotypic plasticity, including acclimation, may provide alternative strategies for surviving increased temperature variation. As mentioned before, tolerant traits may have a more labile response to environmental changes than resistant traits (McNew et al. 2019). Furthermore, mathematical models have found that parasite-tolerant traits increase with environmental variation (Ferris and Best 2019), lending yet another possible avenue of how their evolutionarily different defenses could be shaped by increasingly variable temperatures.

Our findings highlight key areas for future theoretical progress. Notably, we considered only one form of temperature variation over time, as each day had a single temperature drawn from a fixed normal distribution. In reality, temperature varies within days, seasons, and displays additional patterns such as autocorrelation, all of which will alter how temperature shapes epidemics (Gibb et al. 2020; Meadows et al. 2023; Wilber et al. 2022; Donnelly et al. 2013) and thus the relationships between temperature, disease, and host evolution. Additionally, ambient temperatures do not completely dictate the body temperatures of ectotherms, e.g., due to behavioral thermoregulation (Duclaux, Fantino, and Cabanac 1972; Kefford et al. 2022). Overall, we expect that the thermoregulatory abilities of animal ectotherms would effectively reduce temperature variation in most cases. However, intraspecific variation in behavioral thermoregulation (for example, in thermal preference) raises intriguing questions for future theoretical work investigating how such variation within and between populations alters key epidemic and evolutionary outcomes. Lastly, we note that the strong impacts we found of temperature variability on epidemics and host evolution arose from history-independent impacts of temperature, i.e., current epidemiological and demographic rates were determined by current temperatures. However, we know that many relevant rates are history-dependent, meaning previous temperatures, the rates of change, and the time allowed for acclimation play vital roles (Thomas R. Raffel et al. 2013; Rohr et al. 2018; Altman et al. 2016; Noelker et al. 2024). While our model only considers daily temperature variation, incorporating thermal history and temporal autocorrelation (e.g., prolonged temperature events, seasonality, etc.) may more realistically capture the persistent selective pressures that shape evolution. Such history-dependent temperature impacts will yield novel insights into the effects of temperature variability on epidemics and host evolution that we do not currently understand.

As our model focused solely on host evolution, future theoretical and empirical work should combine studies of parasite and host evolution to predict the impacts of temperature variation on disease systems. Parasites and pathogens are often expected to evolve faster than their hosts due to (typically) larger population sizes and faster generation times. Because of the high speed of parasite evolution, previous work has examined how temperature affects parasite evolution (T. E. Hector, Gehman, and King 2023; T. Hector et al. 2025). Given the interactions between temperature and host immune defenses reviewed above, temperature and temperature variability will have strong impacts on outcomes from host evolution (as modeled here), parasite evolution, and host-parasite co-evolution. Understanding these outcomes will require careful measurements of critical, currently untested aspects of exactly how temperature interacts with host immune defenses and their evolution.

In addition to theoretical studies as done here, empirical work is needed that includes monitoring of both resistance and tolerance under temperature variation over eco-evolutionary timescales. Furthermore, how tolerance versus resistance influences host thermal breadth could elucidate potential mechanisms if differences are observed. *Caenorhabditis elegans* serves as an ideal model organism for studying how environmental stressors, particularly temperature variability, interact with host defense strategies. *C. elegans* has known “resistant” and “tolerant” phenotypes to microsporidia or others that could be used to compare the fitness of the host genotypes at varying temperatures with and without infection (Mok et al. 2023; Rafaluk-Mohr et al. 2022). This type of experiment would help establish the ecological context. Furthermore, *C. elegans* has short generation times and could be used to investigate host defense evolution under dual stressors, thereby advancing evolutionary theory and refining predictive models (T. E. Hector, Gehman, and King 2023; Will et al. 2025; Jingdi Li et al. 2024). For other systems with known resistant or tolerant individuals within populations, e.g., House finches and *Mycoplasma* conjunctivitis (Henschen et al. 2023; Kuttiyarthu Veetil et al. 2024) and bullfrogs (*Rana catesbeiana*) and chytridiomycosis caused by Bd (Eskew et al. 2015), we could test the thermal ranges of these phenotypes.

Another missing piece of information needed to fully understand these dynamics is how an individual host’s thermal range is affected long after infection. We know that infection can lower a host’s thermal tolerance, and some evidence suggests that it can recover (Bakewell et al. 2025). However, tracking multiple individuals’ thermal ranges across periods of susceptibility, infection, and recovery will deepen our understanding of how baselines can be affected over time. This could have a profound influence on hosts, favoring those that can limit infection to survive in the long term and under increasing temperature variation, hence, favoring resistance strategies.

While most current research on temperature variability focuses on poikilothermic hosts, the efficiency of the host’s immune system is strongly temperature-dependent (Magnadóttir 2006). However, as climate change progresses and extreme temperature shifts become more common, species across the thermoregulation spectrum will need to be considered. In fact, they likely already should be; human epidemiological studies have shown that temperature variability is associated with increased stroke frequency and mortality among cancer patients (Yi et al. 2021; Chen et al. 2022). Furthermore, in sheep, researchers have found a strong genomic signal associated with isothermality, suggesting that Merino sheep populations may have undergone strong selective pressure due to daily and seasonal temperature variability (Di Civita et al. 2026). Isothermality quantifies the difference between daily temperature fluctuations and the yearly average temperature fluctuation; in other words, it describes how large typical day-to-day temperature swings are relative to the differences between the hottest and coldest months of the year. In addition, it appears that sheep populations differ in infection rates and selection of immune genes. Future research within this system could reveal important genetic differences arising from temperature and parasite infection pressures.

In conclusion, our model and review of the literature suggest that investigating the cornerstone of host defense evolution, parasite load, and temperature variability could prove fruitful in predicting future epidemics and spillover. This interaction between topics has been largely ignored in current research, and we hope to spur further research in this area, particularly as disease outbreaks are on the rise (CITE). Beyond single populations, parasite-tolerant species can expand more rapidly than more sensitive ones and live longer than expected, given their body size (Brook et al. 2023; Guth et al. 2022). In fact, the introduction of known tolerant species has been found to increase parasite prevalence and decrease the host density of less tolerant, sensitive species (Borzée et al. 2017). These findings and hypotheses raise profound questions for future research and the protection of biodiversity and environmental health.

## Acknowledgments

This work was supported by a National Science Foundation (NSF) Biology Integration Institute grant 2120084 to MEBO, which also supported JCW and JMC, an NSF Postdoctoral Research Fellowship in Biology (2305659) to JMC, and a Howard Hughes Medical Institute (Gilliam Fellowship for Advanced Study) to RDK. RDK was also supported by NSF grant 2232985.

## Author contributions

RDK: Investigation, conceptualization, funding acquisition, writing original draft, editing original draft. JCW: Investigation, conceptualization, funding acquisition, writing original draft, editing original draft. JMC: Investigation, funding acquisition, writing original draft, editing original draft. MEBO: Investigation, project administration, conceptualization, funding acquisition, writing original draft, editing original draft.

## References

Altizer, Sonia, Richard S. Ostfeld, Pieter T. J. Johnson, Susan Kutz, and C. Drew Harvell. 2013. “Climate Change and Infectious Diseases: From Evidence to a Predictive Framework.” Science 341 (6145): 514–19.

Altman, Karie A., Sara H. Paull, Pieter T. J. Johnson, Michelle N. Golembieski, Jeffrey P. Stephens, Bryan E. LaFonte, and Thomas R. Raffel. 2016. “Host and Parasite Thermal Acclimation Responses Depend on the Stage of Infection.” Journal of Animal Ecology 85 (4): 1014–24.

Amo, Laura, Hemanta K. Kole, Bethany Scott, Chen-Feng Qi, Juan Wu, and Silvia Bolland. 2021. “CCL17-Producing cDC2s Are Essential in End-Stage Lupus Nephritis and Averted by a Parasitic Infection.” The Journal of Clinical Investigation 131 (11). 10.1172/JCI148000.

Ayres, Janelle S., and David S. Schneider. 2012. “Tolerance of Infections.” Annual Review of Immunology 30 (1): 271–94.

Bakewell, Leah, Carrie Alfonso, Karla A. Alujević, Samantha S. Fontaine, Jaden Keller, Yanileth F. Lopez-Tacoaman, Nathaly E. Ponce-Chilan, et al. 2025. “Higher Parasite Load Is Associated with Lower Heat Tolerance in a Tropical Lizard.” The Journal of Experimental Biology 228 (18): jeb250580.

Best, A., A. White, and M. Boots. 2008. “Maintenance of Host Variation in Tolerance to Pathogens and Parasites.” Proceedings of the National Academy of Sciences of the United States of America 105 (52): 20786–91.

Blanford, Simon, Matthew B. Thomas, Clare Pugh, and Judy K. Pell. 2003. “Temperature Checks the Red Queen? Resistance and Virulence in a Fluctuating Environment: Temperature Checks the Red Queen?” Ecology Letters 6 (1): 2–5.

Bly, J. E., and L. W. Clem. 1991. “Temperature-Mediated Processes in Teleost Immunity: In Vitro Immunosuppression Induced by in Vivo Low Temperature in Channel Catfish.” Veterinary Immunology and Immunopathology 28 (3–4): 365–77.

Bonneaud, Camille, Luc Tardy, Mathieu Giraudeau, Geoffrey E. Hill, Kevin J. McGraw, and Alastair J. Wilson. 2019. “Evolution of Both Host Resistance and Tolerance to an Emerging Bacterial Pathogen.” Evolution Letters 3 (5): 544–54.

Boots, M., and R. G. Bowers. 1999. “Three Mechanisms of Host Resistance to Microparasites-Avoidance, Recovery and Tolerance-Show Different Evolutionary Dynamics.” Journal of Theoretical Biology 201 (1): 13–23.

Borzée, Amaël, Tiffany A. Kosch, Miyeon Kim, and Yikweon Jang. 2017. “Introduced Bullfrogs Are Associated with Increased Batrachochytrium Dendrobatidis Prevalence and Reduced Occurrence of Korean Treefrogs.” PloS One 12 (5): e0177860.

Bozinovic, Francisco, Tamara P. Catalán, and Alexis M. Kalergis. 2013. “Immunological Vulnerability and Adjustments to Environmental Thermal Variability.” Revista Chilena de Historia Natural (Valparaiso, Chile: 1983) 86 (4): 475–83.

Brook, Cara E., Carly Rozins, Sarah Guth, and Mike Boots. 2023. “Reservoir Host Immunology and Life History Shape Virulence Evolution in Zoonotic Viruses.” PLoS Biology 21 (9): e3002268.

Buckley, Lauren B., and Raymond B. Huey. 2016. “Temperature Extremes: Geographic Patterns, Recent Changes, and Implications for Organismal Vulnerabilities.” Global Change Biology 22 (12): 3829–42.

Budischak, Sarah A., and Clayton E. Cressler. 2018. “Fueling Defense: Effects of Resources on the Ecology and Evolution of Tolerance to Parasite Infection.” Frontiers in Immunology 9 (October): 2453.

Butler, Michael W., Zachary R. Stahlschmidt, Daniel R. Ardia, Scott Davies, Jon Davis, Louis J. Guillette Jr, Nicholas Johnson, Stephen D. McCormick, Kevin J. McGraw, and Dale F. DeNardo. 2013. “Thermal Sensitivity of Immune Function: Evidence against a Generalist-Specialist Trade-off among Endothermic and Ectothermic Vertebrates.” The American Naturalist 181 (6): 761–74.

Byrne, Allison Q., Andrew P. Rothstein, Roland A. Knapp, and Erica Bree Rosenblum. 2026. “Genomic Correlates of Disease Recovery in Natural Populations of the Sierra Nevada Yellow-Legged Frog (Rana Sierrae).” Molecular Ecology 35 (8): e70348.

Cao, Chang, Vishwajit S. Chowdhury, Mark A. Cline, and Elizabeth R. Gilbert. 2021. “The Microbiota-Gut-Brain Axis during Heat Stress in Chickens: A Review.” Frontiers in Physiology 12 (October): 752265.

Chen, Zhuangzhuang, Peilin Liu, Xiaoshuang Xia, Lin Wang, and Xin Li. 2022. “The Underlying Mechanisms of Cold Exposure-Induced Ischemic Stroke.” The Science of the Total Environment 834 (155514): 155514.

Clusella-Trullas, Susana, Tim M. Blackburn, and Steven L. Chown. 2011. “Climatic Predictors of Temperature Performance Curve Parameters in Ectotherms Imply Complex Responses to Climate Change.” The American Naturalist 177 (6): 738–51.

Cohen, Jeremy M., Matthew D. Venesky, Erin L. Sauer, David J. Civitello, Taegan A. McMahon, Elizabeth A. Roznik, and Jason R. Rohr. 2017. “The Thermal Mismatch Hypothesis Explains Host Susceptibility to an Emerging Infectious Disease.” Ecology Letters 20 (2): 184–93.

Debes, Paul Vincent, Riho Gross, and Anti Vasemägi. 2017. “Quantitative Genetic Variation in, and Environmental Effects on, Pathogen Resistance and Temperature-Dependent Disease Severity in a Wild Trout.” The American Naturalist 190 (2): 244–65.

Deutsch, Curtis A., Joshua J. Tewksbury, Raymond B. Huey, Kimberly S. Sheldon, Cameron K. Ghalambor, David C. Haak, and Paul R. Martin. 2008. “Impacts of Climate Warming on Terrestrial Ectotherms across Latitude.” Proceedings of the National Academy of Sciences of the United States of America 105 (18): 6668–72.

Di Civita, M., P. Wiener, M. Marr, C. Persichilli, G. Senczuk, F. Pilla, E. L. Clark, and J. Friedrich. 2026. “Exploring Climate Adaptation in European Merino Sheep: A Landscape Genomics Approach.” Animal: An International Journal of Animal Bioscience 20 (1): 101728.

Doeschl-Wilson, Andrea B., Will Brindle, Gerry Emmans, and Ilias Kyriazakis. 2009. “Unravelling the Relationship between Animal Growth and Immune Response during Micro-Parasitic Infections.” PloS One 4 (10): e7508.

Donnelly, R., A. Best, A. White, and M. Boots. 2013. “Seasonality Selects for More Acutely Virulent Parasites When Virulence Is Density Dependent.” *Proceedings*. Biological Sciences 280 (1751): 20122464.

Duclaux, R., M. Fantino, and M. Cabanac. 1972. “Thermoregulatory Behavior of Frogs - Influence of Spinal Thermal Stimulation.” Journal de Physiologie 65 (3): A392–A392.

Duncan, A. B., E. Dusi, F. Jacob, J. Ramsayer, M. E. Hochberg, and O. Kaltz. 2017. “Hot Spots Become Cold Spots: Coevolution in Variable Temperature Environments.” Journal of Evolutionary Biology 30 (1): 55–65.

Easterling, D. R., G. A. Meehl, C. Parmesan, S. A. Changnon, T. R. Karl, and L. O. Mearns. 2000. “Climate Extremes: Observations, Modeling, and Impacts.” Science (New York, N.Y.) 289 (5487): 2068–74.

Eskew, Evan A., S. Joy Worth, Janet E. Foley, and Brian D. Todd. 2015. “American Bullfrogs (Lithobates Catesbeianus) Resist Infection by Multiple Isolates of Batrachochytrium Dendrobatidis, Including One Implicated in Wild Mass Mortality.” EcoHealth 12 (3): 513–18.

Ferris, Charlotte, and Alex Best. 2018. “The Evolution of Host Defence to Parasitism in Fluctuating Environments.” Journal of Theoretical Biology 440 (March): 58–65.

Ferris, Charlotte, and Alex Best. 2019. “The Effect of Temporal Fluctuations on the Evolution of Host Tolerance to Parasitism.” Theoretical Population Biology 130 (December): 182–90.

Fontaine, Samantha S., and Kevin D. Kohl. 2023. “The Microbiome Buffers Tadpole Hosts from Heat Stress: A Hologenomic Approach to Understand Host-Microbe Interactions under Warming.” The Journal of Experimental Biology 226 (1). 10.1242/jeb.245191.

Gehman, Alyssa-Lois M., Richard J. Hall, and James E. Byers. 2018. “Host and Parasite Thermal Ecology Jointly Determine the Effect of Climate Warming on Epidemic Dynamics.” Proceedings of the National Academy of Sciences of the United States of America 115 (4): 744–49.

Gibb, Rory, Lydia H. V. Franklinos, David W. Redding, and Kate E. Jones. 2020. “Ecosystem Perspectives Are Needed to Manage Zoonotic Risks in a Changing Climate.” BMJ (Clinical Research Ed.) 371 (November): m3389.

Grant, Peter R., B. Rosemary Grant, Raymond B. Huey, Marc T. J. Johnson, Andrew H. Knoll, and Johanna Schmitt. 2017. “Evolution Caused by Extreme Events.” Philosophical Transactions of the Royal Society of London. Series B, Biological Sciences 372 (1723). 10.1098/rstb.2016.0146.

Greenspan, Sasha E., Deborah S. Bower, Elizabeth A. Roznik, David A. Pike, Gerry Marantelli, Ross A. Alford, Lin Schwarzkopf, and Brett R. Scheffers. 2017. “Infection Increases Vulnerability to Climate Change via Effects on Host Thermal Tolerance.” Scientific Reports 7 (1): 9349.

Greenspan, Sasha E., Deborah S. Bower, Rebecca J. Webb, Lee Berger, Donna Rudd, Lin Schwarzkopf, and Ross A. Alford. 2017. “White Blood Cell Profiles in Amphibians Help to Explain Disease Susceptibility Following Temperature Shifts.” Developmental and Comparative Immunology 77 (December): 280–86.

Guth, Sarah, Nardus Mollentze, Katia Renault, Daniel G. Streicker, Elisa Visher, Mike Boots, and Cara E. Brook. 2022. “Bats Host the Most Virulent-but Not the Most Dangerous-Zoonotic Viruses.” Proceedings of the National Academy of Sciences of the United States of America 119 (14): e2113628119.

Gutschick, Vincent P., and Hormoz BassiriRad. 2003. “Extreme Events as Shaping Physiology, Ecology, and Evolution of Plants: Toward a Unified Definition and Evaluation of Their Consequences: Tansley Review.” The New Phytologist 160 (1): 21–42.

Handel A., Rohani P. 2015. “Crossing the Scale from Within-Host Infection Dynamics to between-Host Transmission Fitness: A Discussion of Current Assumptions and Knowledge.” Philosophical Transactions of the Royal Society B: Biological Sciences 370. https://www.ncbi.nlm.nih.gov/pmc/articles/PMC4528500/.

Harrison, Ellie, Anna-Liisa Laine, Mikael Hietala, and Michael A. Brockhurst. 2013. “Rapidly Fluctuating Environments Constrain Coevolutionary Arms Races by Impeding Selective Sweeps.” *Proceedings*. Biological Sciences 280 (1764): 20130937.

Hayward, Adam D., Romain Garnier, Kathryn A. Watt, Jill G. Pilkington, Bryan T. Grenfell, Jacqueline B. Matthews, Josephine M. Pemberton, Daniel H. Nussey, and Andrea L. Graham. 2014. “Heritable, Heterogeneous, and Costly Resistance of Sheep against Nematodes and Potential Feedbacks to Epidemiological Dynamics.” The American Naturalist 184 Suppl 1 (S1): S58–76.

Hector, T. E., A-L M. Gehman, and K. C. King. 2023. “Infection Burdens and Virulence under Heat Stress: Ecological and Evolutionary Considerations.” Philosophical Transactions of the Royal Society of London. Series B, Biological Sciences 378 (1873): 20220018.

Hector, Tobias E., Carla M. Sgrò, and Matthew D. Hall. 2021. “Thermal Limits in the Face of Infectious Disease: How Important Are Pathogens?” Global Change Biology 27 (19): 4469–80.

Hector, Tobias, Julia Kreiner, James Forward, Kim Hoang, Emily Stevens, Serena Johnson, Jingdi Li, and Kayla King. 2025. “Rising Temperatures Favour Parasite Virulence and Parallel Molecular Evolution Following a Host Jump.” bioRxiv. 10.1101/2025.01.14.632940.

Heled, Yuval, Chen Fleischmann, and Yoram Epstein. 2013. “Cytokines and Their Role in Hyperthermia and Heat Stroke.” Journal of Basic and Clinical Physiology and Pharmacology 24 (2): 85–96.

Henschen, Amberleigh E., Michal Vinkler, Marissa M. Langager, Allison A. Rowley, Rami A. Dalloul, Dana M. Hawley, and James S. Adelman. 2023. “Rapid Adaptation to a Novel Pathogen through Disease Tolerance in a Wild Songbird.” PLoS Pathogens 19 (6): e1011408.

Holand, H., H. Jensen, T. Kvalnes, J. Tufto, H. Pärn, B-E Sæther, and T. H. Ringsby. 2019. “Parasite Prevalence Increases with Temperature in an Avian Metapopulation in Northern Norway.” Parasitology 146 (8): 1030–35.

Huntley, John Warren, Franz T. Fürsich, Matthias Alberti, Manja Hethke, and Chunlian Liu. 2014. “A Complete Holocene Record of Trematode-Bivalve Infection and Implications for the Response of Parasitism to Climate Change.” Proceedings of the National Academy of Sciences of the United States of America 111 (51): 18150–55.

Kanjanapruthipong, J., W. Junlapho, and K. Karnjanasirm. 2015. “Feeding and Lying Behavior of Heat-Stressed Early Lactation Cows Fed Low Fiber Diets Containing Roughage and Nonforage Fiber Sources.” Journal of Dairy Science 98 (2): 1110–18.

Kefford, Ben J., Cameron K. Ghalambor, Beatrice Dewenter, N. Leroy Poff, Jane Hughes, Jollene Reich, and Ross Thompson. 2022. “Acute, Diel, and Annual Temperature Variability and the Thermal Biology of Ectotherms.” Global Change Biology 28 (23): 6872–88.

Kirk, Devin, Mary I. O’Connor, and Erin A. Mordecai. 2022. “Scaling Effects of Temperature on Parasitism from Individuals to Populations.” The Journal of Animal Ecology 91 (10): 2087–2102.

Klemme, Ines, and Anssi Karvonen. 2017. “Vertebrate Defense against Parasites: Interactions between Avoidance, Resistance, and Tolerance.” Ecology and Evolution 7 (2): 561–71.

Knapp, Roland A., Gary M. Fellers, Patrick M. Kleeman, David A. W. Miller, Vance T. Vredenburg, Erica Bree Rosenblum, and Cheryl J. Briggs. 2016. “Large-Scale Recovery of an Endangered Amphibian despite Ongoing Exposure to Multiple Stressors.” Proceedings of the National Academy of Sciences of the United States of America 113 (42): 11889–94.

Knapp, Roland A., Mark Q. Wilber, Maxwell B. Joseph, Thomas C. Smith, and Robert L. Grasso. 2024. “Reintroduction of Resistant Frogs Facilitates Landscape-Scale Recovery in the Presence of a Lethal Fungal Disease.” Nature Communications 15 (1): 9436.

Knutie, Sarah A., Christina L. Wilkinson, Qiu Chang Wu, C. Nicole Ortega, and Jason R. Rohr. 2017. “Host Resistance and Tolerance of Parasitic Gut Worms Depend on Resource Availability.” Oecologia 183 (4): 1031–40.

Koelle, Katia, Mercedes Pascual, and Md Yunus. 2005. “Pathogen Adaptation to Seasonal Forcing and Climate Change.” *Proceedings*. Biological Sciences 272 (1566): 971–77.

Kramp, Rachael D., Mary J. Janecka, Nadine Tardent, Jukka Jokela, Kevin D. Kohl, and Jessica F. Stephenson. 2026. “Host Genetics and the Skin Microbiome Independently Predict Parasite Resistance.” Ecology and Evolution 16 (1): e72923.

Krichel, Leila, Devin Kirk, Clara Pencer, Madison Hönig, Kiran Wadhawan, and Martin Krkošek. 2023. “Short-Term Temperature Fluctuations Increase Disease in a Daphnia-Parasite Infectious Disease System.” PLoS Biology 21 (9): e3002260.

Kuttiyarthu Veetil, Nithya, Amberleigh E. Henschen, Dana M. Hawley, Balraj Melepat, Rami A. Dalloul, Vladimír Beneš, James S. Adelman, and Michal Vinkler. 2024. “Varying Conjunctival Immune Response Adaptations of House Finch Populations to a Rapidly Evolving Bacterial Pathogen.” Frontiers in Immunology 15 (February): 1250818.

Kyprianou, Themis-Dimitrios, Hans O. Pörtner, Andreas Anestis, Basile Kostoglou, Konstantinos Feidantsis, and Basile Michaelidis. 2010. “Metabolic and Molecular Stress Responses of Gilthead Seam Bream Sparus Aurata during Exposure to Low Ambient Temperature: An Analysis of Mechanisms Underlying the Winter Syndrome.” Journal of Comparative Physiology. B, Biochemical, Systemic, and Environmental Physiology 180 (7): 1005–18.

Li, J. D., Y. Y. Gao, E. J. Stevens, and K. C. King. 2024. “Dual Stressors of Infection and Warming Can Destabilize Host Microbiomes.” Philosophical Transactions of the Royal Society of London. Series B, Biological Sciences 379 (1901): 20230069.

Li, Jiaying, Jiahao Zhu, Ayi Budian, Yiwei Zeng, Shouhong Wang, Yuzhou Gong, Xungang Wang, et al. 2025. “Metabolic Acclimation to Warming Links Unexpected Immune Activation and Sexual Dimorphism Attenuation in Xenopus Tropicalis.” Communications Biology 8 (1): 952.

Li, Jingdi, Cameron Smith, Jinlin Chen, and Kayla King. 2024. “Warming during Different Life Stages Has Distinct Impacts on Host Resistance Ecology and Evolution.” Authorea Inc. 10.22541/au.172457643.32930358/v1.

Linderman, Jessica A., Moria C. Chambers, Avni S. Gupta, and David S. Schneider. 2012. “Infection-Related Declines in Chill Coma Recovery and Negative Geotaxis in Drosophila Melanogaster.” PloS One 7 (9): e41907.

Liu, Elaine, Kevin Lewis, Hiba Al-Saffar, Catherine M. Krall, Anju Singh, Vladimir A. Kulchitsky, Joshua J. Corrigan, et al. 2012. “Naturally Occurring Hypothermia Is More Advantageous than Fever in Severe Forms of Lipopolysaccharide- and Escherichia Coli-Induced Systemic Inflammation.” American Journal of Physiology. Regulatory, Integrative and Comparative Physiology 302 (12): R1372–83.

Liu, Qi, Congbin Fu, Zhongfeng Xu, and Aijun Ding. 2026. “Global Warming Intensifies Extreme Day-to-Day Temperature Changes in Mid–Low Latitudes.” Nature Climate Change 16 (1): 69–76.

Luong, L. T., and M. Polak. 2007. “Environment-Dependent Trade-Offs between Ectoparasite Resistance and Larval Competitive Ability in the Drosophila-Macrocheles System.” Heredity 99 (6): 632–40.

Magnadóttir, Bergljót. 2006. “Innate Immunity of Fish (Overview).” Fish & Shellfish Immunology 20 (2): 137–51.

McCartan, Niamh, Floriane O’Keeffe, Guoyuan Zhang, and Pepijn Luijckx. 2025. “Impact of Heatwave Amplitude, Duration, and Timing on Parasite Fitness at Different Baseline Temperatures.” PLOS Climate 4 (6): e0000632.

McCartan, Niamh, Jeremy Piggott, Sadie DiCarlo, and Pepijn Luijckx. 2024. “Cold Snaps Lead to a 5-Fold Increase or a 3-Fold Decrease in Disease Proliferation Depending on the Baseline Temperature.” BMC Biology 22 (1): 250.

McCarville, J. L., and J. S. Ayres. 2018. “Disease Tolerance: Concept and Mechanisms.” Current Opinion in Immunology 50 (February): 88–93.

McNew, Sabrina M., Sarah A. Knutie, Graham B. Goodman, Angela Theodosopoulos, Ashley Saulsberry, Janai Yépez R, Sarah E. Bush, and Dale H. Clayton. 2019. “Annual Environmental Variation Influences Host Tolerance to Parasites.” Proceedings. Biological Sciences 286 (1897): 20190049.

Meadows, Amanda Jean, Nicole Stephenson, Nita K. Madhav, and Ben Oppenheim. 2023. “Historical Trends Demonstrate a Pattern of Increasingly Frequent and Severe Spillover Events of High-Consequence Zoonotic Viruses.” BMJ Global Health 8 (11): e012026.

Meng, Di, Yanxin Hu, Chong Xiao, Tangting Wei, Qiang Zou, and Ming Wang. 2013. “Chronic Heat Stress Inhibits Immune Responses to H5N1 Vaccination through Regulating CD4^+^ CD25^+^ Foxp3^+^ Tregs.” BioMed Research International 2013 (September): 160859.

Mideo, Nicole, Samuel Alizon, and Troy Day. 2008. “Linking Within- and between-Host Dynamics in the Evolutionary Epidemiology of Infectious Diseases.” Trends in Ecology & Evolution 23 (9): 511–17.

Miller, M. R., A. White, and M. Boots. 2005. “The Evolution of Host Resistance: Tolerance and Control as Distinct Strategies.” Journal of Theoretical Biology 236 (2): 198–207.

Miller, Martin R., Andrew White, and Michael Boots. 2006. “The Evolution of Parasites in Response to Tolerance in Their Hosts: The Good, the Bad, and Apparent Commensalism.” Evolution; International Journal of Organic Evolution 60 (5): 945–56.

Mok, Calvin, Meng A. Xiao, Yin C. Wan, Winnie Zhao, Shanzeh M. Ahmed, Robert J. Luallen, and Aaron W. Reinke. 2023. “High-Throughput Phenotyping of Infection by Diverse Microsporidia Species Reveals a Wild C. Elegans Strain with Opposing Resistance and Susceptibility Traits.” PLoS Pathogens 19 (3): e1011225.

Noelker, James E., Vitoria Abreu Ruozzi, Kyle D. Spengler, Hunter M. Craig, and Thomas R. Raffel. 2024. “Dynamic Effects of Thermal Acclimation on Chytridiomycosis Infection Intensity and Transmission Potential in Xenopus Laevis.” Royal Society Open Science 11 (9): 240789.

Paaijmans, Krijn P., Rebecca L. Heinig, Rebecca A. Seliga, Justine I. Blanford, Simon Blanford, Courtney C. Murdock, and Matthew B. Thomas. 2013. “Temperature Variation Makes Ectotherms More Sensitive to Climate Change.” Global Change Biology 19 (8): 2373–80.

Patel, Ronak N., David B. Bonan, and Tapio Schneider. 2024. “Changes in the Frequency of Observed Temperature Extremes Largely Driven by a Distribution Shift.” Geophysical Research Letters 51 (24). 10.1029/2024gl110707.

Pisano, Olivia M., Anna Kuparinen, and Jeffrey A. Hutchings. 2019. “Cyclical and Stochastic Thermal Variability Affects Survival and Growth in Brook Trout.” Journal of Thermal Biology, Cvequality: Tests for the Equality of Coefficients of Variation from Multiple Groups. R Software Package Version 0.1.3R: A Language and Environment for Statistical ComputingModelling Survival Data: Extending the Cox Model, 84 (August): 221–27.

Pottier, Patrice, Rachel R. Y. Oh, Pietro Pollo, A. Nayelli Rivera-Villanueva, Yefeng Yang, Sarah Varon, Ana V. Longo, et al. 2025. “AmphiTherm: A Comprehensive Database of Amphibian Thermal Tolerance and Preference.” Scientific Data 12 (1): 1987.

Poulton, Anna J., and Stephen P. Ellner. 2025. “Learned Behavioral Avoidance Can Alter Outbreak Dynamics in a Model for Waterborne Infectious Diseases.” Journal of Mathematical Biology 91 (3): 28.

Quance, Michael A., and Michael Travisano. 2009. “Effects of Temperature on the Fitness Cost of Resistance to Bacteriophage T4 in Escherichia Coli.” Evolution; International Journal of Organic Evolution 63 (6): 1406–16.

Råberg, Lars. 2014. “How to Live with the Enemy: Understanding Tolerance to Parasites.” PLoS Biology 12 (11): e1001989.

Råberg, Lars, Andrea L. Graham, and Andrew F. Read. 2009. “Decomposing Health: Tolerance and Resistance to Parasites in Animals.” Philosophical Transactions of the Royal Society of London. Series B, Biological Sciences 364 (1513): 37–49.

Råberg, Lars, Derek Sim, and Andrew F. Read. 2007. “Disentangling Genetic Variation for Resistance and Tolerance to Infectious Diseases in Animals.” Science 318 (5851): 812–14.

Rafaluk-Mohr, Charlotte, Michael Gerth, Jordan E. Sealey, Alice K. E. Ekroth, Aziz A. Aboobaker, Anke Kloock, and Kayla C. King. 2022. “Microbial Protection Favors Parasite Tolerance and Alters Host-Parasite Coevolutionary Dynamics.” *Current Biology: CB*, February. 10.1016/j.cub.2022.01.063.

Raffel, T. R., J. R. Rohr, J. M. Kiesecker, and P. J. Hudson. 2006. “Negative Effects of Changing Temperature on Amphibian Immunity under Field Conditions.” Functional Ecology 20 (5): 819–28.

Raffel, Thomas R., Neal T. Halstead, Taegan A. McMahon, Andrew K. Davis, and Jason R. Rohr. 2015. “Temperature Variability and Moisture Synergistically Interact to Exacerbate an Epizootic Disease.” *Proceedings*. Biological Sciences 282 (1801): 20142039.

Raffel, Thomas R., John M. Romansic, Neal T. Halstead, Taegan A. McMahon, Matthew D. Venesky, and Jason R. Rohr. 2013. “Disease and Thermal Acclimation in a More Variable and Unpredictable Climate.” Nature Climate Change 3 (2): 146–51.

Rebl, Alexander, Marieke Verleih, Mareen Nipkow, Simone Altmann, Ralf Bochert, and Tom Goldammer. 2018. “Gradual and Acute Temperature Rise Induces Crossing Endocrine, Metabolic, and Immunological Pathways in Maraena Whitefish (Coregonus Maraena).” Frontiers in Genetics 9 (July): 241.

Restif, Olivier, and Jacob C. Koella. 2004. “Concurrent Evolution of Resistance and Tolerance to Pathogens.” The American Naturalist 164 (4): E90–102.

Rijkers, G. T., E. M. Frederix-Wolters, and W. B. van Muiswinkel. 1980. “The Immune System of Cyprinid Fish. Kinetics and Temperature Dependence of Antibody-Producing Cells in Carp (Cyprinus Carpio).” Immunology 41 (1): 91–97.

Rijkers, G. T., J. A. M. Wiegerinck, R. van Oosterom, and W. B. van Muiswinkel. 1981. “Temperature Dependence of Humoral Immunity in Carp (Cyprinus Carpio).” In Aspects of Developmental and Comparative Immunology, 477–82. Elsevier.

Rohr, Jason R., David J. Civitello, Jeremy M. Cohen, Elizabeth A. Roznik, Barry Sinervo, and Anthony I. Dell. 2018. “The Complex Drivers of Thermal Acclimation and Breadth in Ectotherms.” Ecology Letters 21 (9): 1425–39.

Rohr, Jason R., and Jeremy M. Cohen. 2020. “Understanding How Temperature Shifts Could Impact Infectious Disease.” PLoS Biology 18 (11): e3000938.

Rosa, Gonçalo M., Rachel Perez, Lora A. Richards, Corinne L. Richards-Zawacki, Angela M. Smilanich, Laura K. Reinert, Louise A. Rollins-Smith, Daniel P. Wetzel, and Jamie Voyles. 2022. “Seasonality of Host Immunity in a Tropical Disease System.” Ecosphere (Washington, D.C) 13 (7): e4158.

Roy, B. A., and J. W. Kirchner. 2000. “Evolutionary Dynamics of Pathogen Resistance and Tolerance.” Evolution; International Journal of Organic Evolution 54 (1): 51–63.

Saxon, A. D., E. K. O’Brien, and J. R. Bridle. 2018. “Temperature Fluctuations during Development Reduce Male Fitness and May Limit Adaptive Potential in Tropical Rainforest Drosophila.” Journal of Evolutionary Biology 31 (3): 405–15.

Scanes, Elliot, Nachshon Siboni, Brendon Rees, and Justin R. Seymour. 2023. “Acclimation in Intertidal Animals Reduces Potential Pathogen Load and Increases Survival Following a Heatwave.” iScience 26 (6): 106813.

Scharsack, Jörn P., Bartholomäus Wieczorek, Alexander Schmidt-Drewello, Janine Büscher, Frederik Franke, Andrew Moore, Antoine Branca, et al. 2021. “Climate Change Facilitates a Parasite’s Host Exploitation via Temperature-Mediated Immunometabolic Processes.” Global Change Biology 27 (1): 94–107.

Scharsack, Jörn Peter, and Frederik Franke. 2022. “Temperature Effects on Teleost Immunity in the Light of Climate Change.” Journal of Fish Biology 101 (4): 780–96.

Schieber, Alexandria M. Palaferri, and Janelle S. Ayres. 2016. “Thermoregulation as a Disease Tolerance Defense Strategy.” Pathogens and Disease 74 (9): ftw106.

Seal, Srijan, Guha Dharmarajan, and Imroze Khan. 2021. “Evolution of Pathogen Tolerance and Emerging Infections: A Missing Experimental Paradigm.” eLife 10 (September). 10.7554/eLife.68874.

Sears, Brittany F., Paul W. Snyder, and Jason R. Rohr. 2015. “Host Life History and Host-Parasite Syntopy Predict Behavioural Resistance and Tolerance of Parasites.” The Journal of Animal Ecology 84 (3): 625–36.

Shakhar, Keren. 2019. “The Inclusive Behavioral Immune System.” Frontiers in Psychology 10 (May): 1004.

Singh, Prerna, and Alex Best. 2021. “Simultaneous Evolution of Host Resistance and Tolerance to Parasitism.” Journal of Evolutionary Biology 34 (12): 1932–43.

Sorrell, Ian, Andrew White, Amy B. Pedersen, Rosemary S. Hails, and Mike Boots. 2009. “The Evolution of Covert, Silent Infection as a Parasite Strategy.” *Proceedings*. Biological Sciences 276 (1665): 2217–26.

Sternberg, Eleanore D., Thierry Lefèvre, James Li, Carlos Lopez Fernandez de Castillejo, Hui Li, Mark D. Hunter, and Jacobus C. de Roode. 2012. “Food Plant Derived Disease Tolerance and Resistance in a Natural Butterfly-Plant-Parasite Interactions: Food Plant-Derived Disease Tolerance and Resistance.” Evolution; International Journal of Organic Evolution 66 (11): 3367–76.

Streipert, Sabrina H., David Swigon, Mark Q. Wilber, and Jason C. Walsman. 2025. “Evolution of Pathogen Tolerance and Reproductive Trade-off Implications.” Journal of Mathematical Biology 90 (5): 53.

Terrell, Kimberly A., Richard P. Quintero, Veronica Acosta Galicia, Ed Bronikowski, Matthew Evans, John D. Kleopfer, Suzan Murray, James B. Murphy, Bradley D. Nissen, and Brian Gratwicke. 2021. “Physiological Impacts of Temperature Variability and Climate Warming in Hellbenders (Cryptobranchus Alleganiensis).” Conservation Physiology 9 (1): coab079.

Terrell, Kimberly A., Richard P. Quintero, Suzan Murray, John D. Kleopfer, James B. Murphy, Matthew J. Evans, Bradley D. Nissen, and Brian Gratwicke. 2013. “Cryptic Impacts of Temperature Variability on Amphibian Immune Function.” The Journal of Experimental Biology 216 (Pt 22): 4204–11.

Ummenhofer, Caroline C., and Gerald A. Meehl. 2017. “Extreme Weather and Climate Events with Ecological Relevance: A Review.” Philosophical Transactions of the Royal Society of London. Series B, Biological Sciences 372 (1723). 10.1098/rstb.2016.0135.

Vasseur, David A., John P. DeLong, Benjamin Gilbert, Hamish S. Greig, Christopher D. G. Harley, Kevin S. McCann, Van Savage, Tyler D. Tunney, and Mary I. O’Connor. 2014. “Increased Temperature Variation Poses a Greater Risk to Species than Climate Warming.” *Proceedings*. Biological Sciences 281 (1779): 20132612.

Walsman, Jason C., Meghan A. Duffy, Carla E. Cáceres, and Spencer R. Hall. 2023. “‘Resistance Is Futile’: Weaker Selection for Resistance by Abundant Parasites Increases Prevalence and Depresses Host Density.” The American Naturalist 201 (6): 864–79.

Walsman, Jason Cosens, Sabrina H. Streipert, Cheryl J. Briggs, and Mark Q. Wilber. 2026. “Dynamic Mean and Variance of Microparasite Load Give Key Insights into Population Dynamics and Underlying Mechanisms.” *Journal of the Royal Society*, Interface 23 (235). 10.1098/rsif.2025.0725.

Wilber, Mark Q., Kate E. Langwig, A. Marm Kilpatrick, Hamish I. McCallum, and Cheryl J. Briggs. 2016. “Integral Projection Models for Host-Parasite Systems with an Application to Amphibian Chytrid Fungus.” Methods in Ecology and Evolution 7 (10): 1182–94.

Wilber, Mark Q., Michel E. B. Ohmer, Karie A. Altman, Laura A. Brannelly, Brandon C. LaBumbard, Emily H. Le Sage, Nina B. McDonnell, et al. 2022. “Once a Reservoir, Always a Reservoir? Seasonality Affects the Pathogen Maintenance Potential of Amphibian Hosts.” Ecology 103 (9): e3759.

Will, I., C. A. Smith, T. E. Hector, and K. C. King. 2025. “Coinfections Dampen the Effects of Temperature on Host-Parasite Interactions.” bioRxiv. bioRxiv. 10.64898/2025.12.10.693475.

Yeh, Sang-Wook, Jong-Seong Kug, Boris Dewitte, Min-Ho Kwon, Ben P. Kirtman, and Fei-Fei Jin. 2009. “El Niño in a Changing Climate.” Nature 461 (7263): 511–14.

Yi, Weizhuo, Jian Cheng, Qiannan Wei, Rubing Pan, Shasha Song, Yangyang He, Chao Tang, Xiangguo Liu, Yu Zhou, and Hong Su. 2021. “Disparities of Weather Type and Geographical Location in the Impacts of Temperature Variability on Cancer Mortality: A Multicity Case-Crossover Study in Jiangsu Province, China.” Environmental Research 197 (110985): 110985.

