## Supplemental information for "Thermal variability interacts with infection load and the evolution of host-parasite defense"

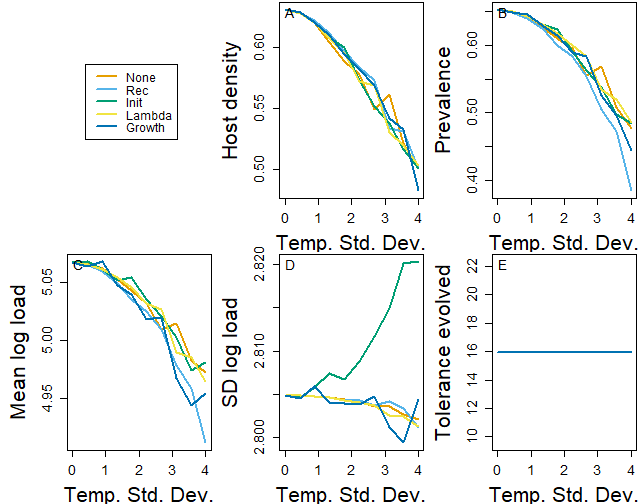

**Figure S1.** Thermal variability depresses (A) host density, (B) prevalence, (C) mean log load, (D) and the standard deviation of log load in most cases, (E) when hosts do not evolve.

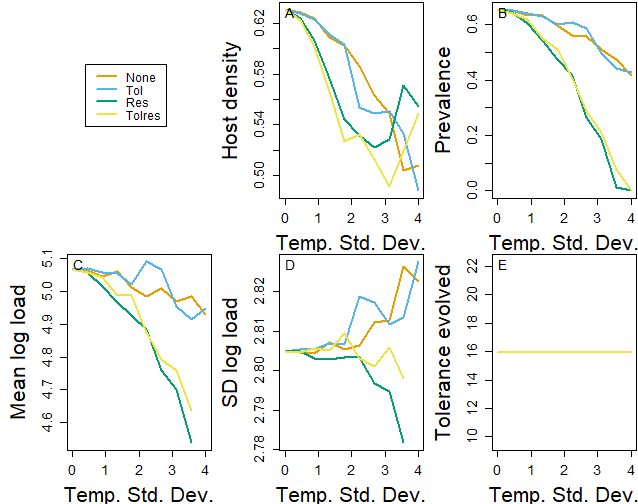

**Figure S2.** Thermal variability depresses (A) host density, (B) prevalence, (C) mean log load, (D) and has mixed effects on the standard deviation of log load, (E) when hosts do not evolve.

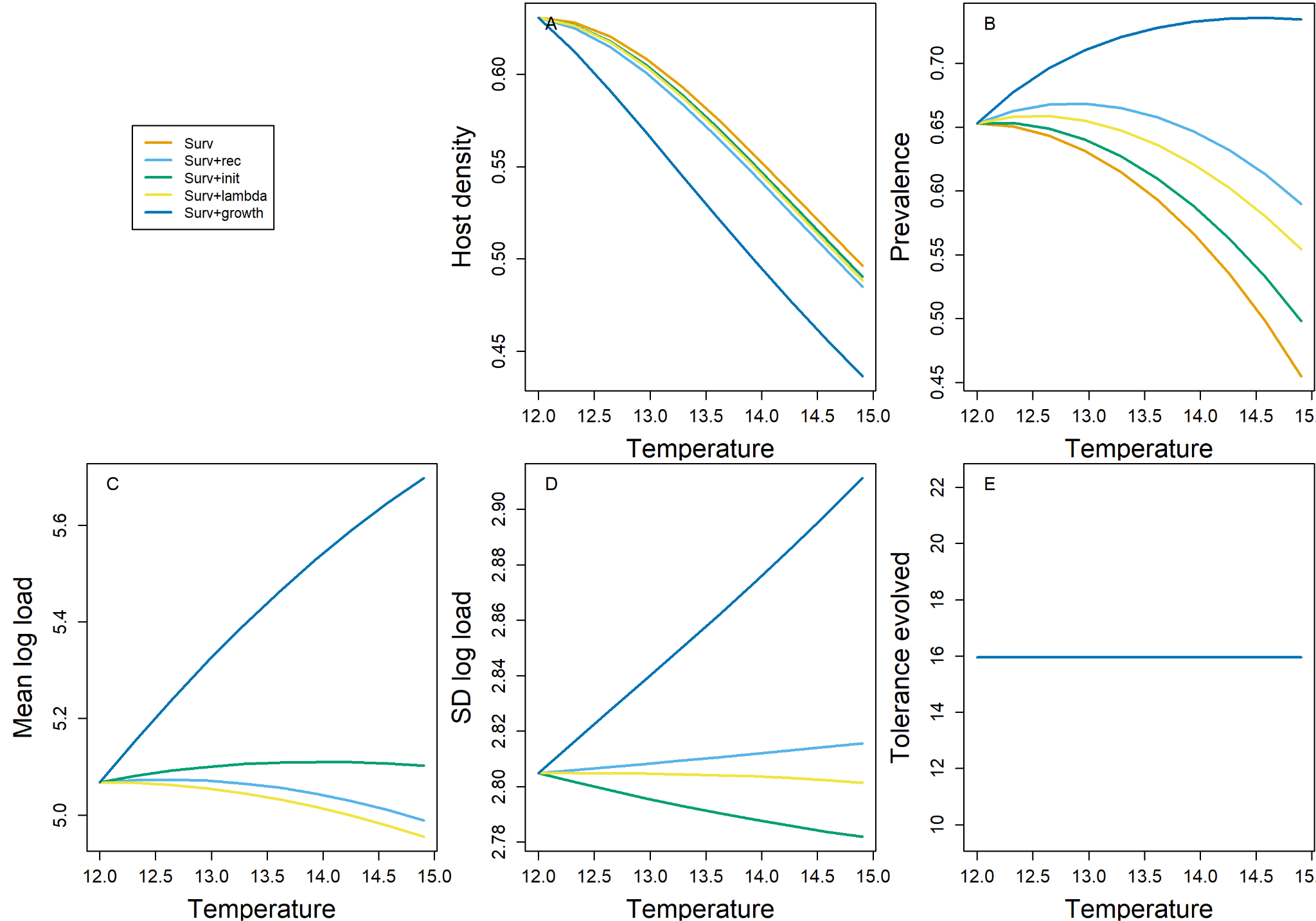

**Figure S3.** Temperature increase depresses (A) host density and has trait-dependent impacts on (B) prevalence, (C) mean log load, (D) and the standard deviation of log load, (E) when hosts do not evolve.

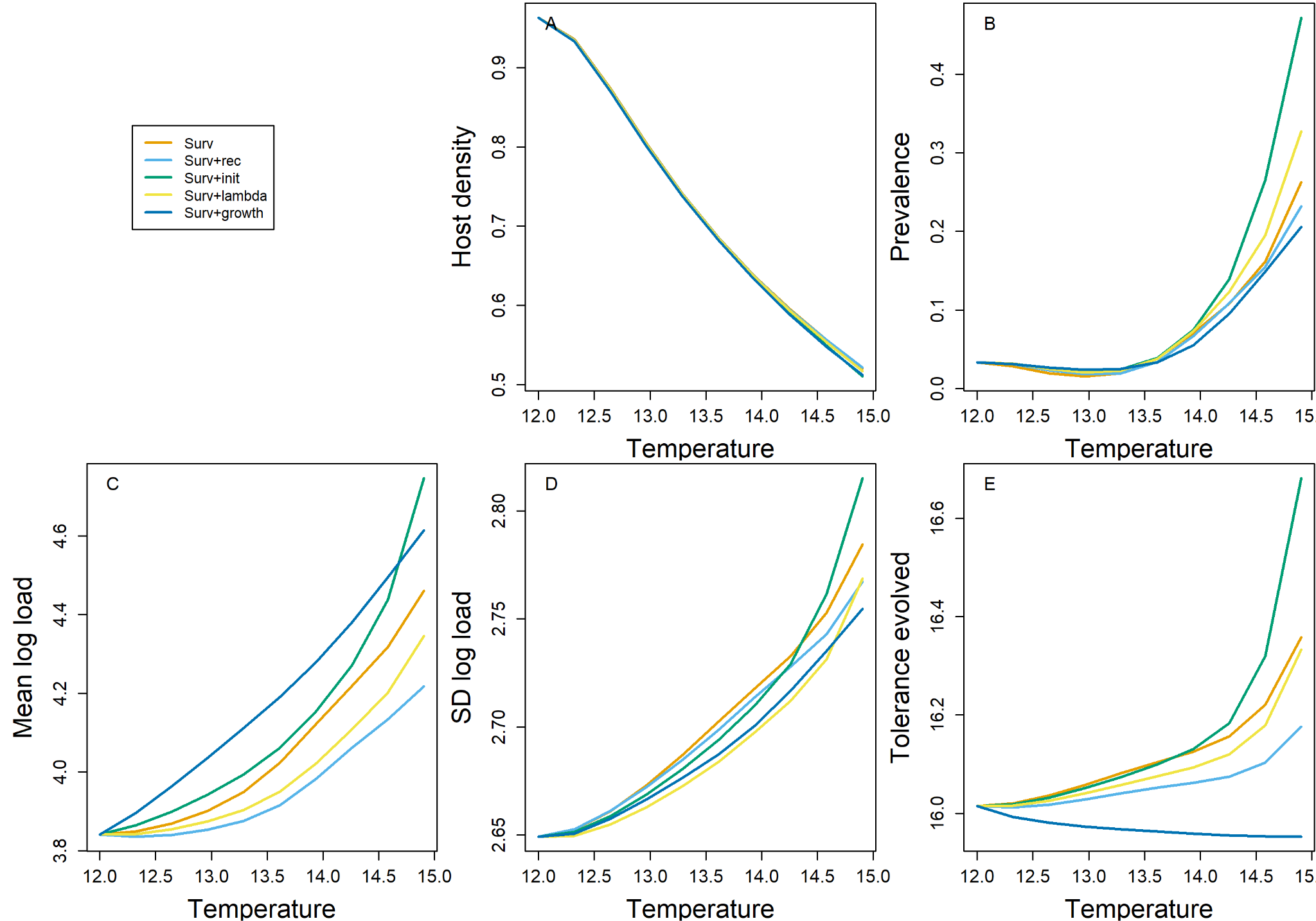

**Figure S4.** Temperature increase depresses (A) host density, (B) increases prevalence, (C) increases mean log load, (D) and increases the standard deviation of log load in most cases, (E) when hosts evolve.

**Table 1.** Overview of factors influencing resistant and tolerant evolution outside of temperature and temperature variability

| **Factor** | **Resistance** | **Tolerance** |
| --- | --- | --- |
| *Infection dynamics* | When a novel pathogen enters a naive population, if the transmission risk and the cost of infection (the parasite's virulence) are high, we should expect that resistance provides optimal protection while maintaining host fitness. Resistance reduces the risk of infection at the individual level and maintains fecundity by reducing the likelihood of infection and preserving the ability to attract mates. Behavioral immunity, a form of resistance, is a rapid behavioral response of avoiding an infected conspecific (Shakhar 2019). Naive hosts can quickly learn to avoid infection and thereby change transmission dynamics (Poulton and Ellner 2025). Within sexually reproducing populations, an individual with the ability to suppress or avoid infection should be able to attract mates more readily than an infected counterpart. | However, if parasite prevalence is high and infection becomes unavoidable, resistance may not be the optimal strategy (Walsman et al. 2023). This is where tolerance may be more beneficial for the host, as it reduces the inevitable damage caused by infection (Budischak and Cressler 2018). Furthermore, the host resistance-tolerance trade-off appears, in some circumstances, to depend on the parasite's lifespan (Balard et al., 2020). |
| *Defence cost* | Both resistance and tolerance come at a cost, and these costs shape which strategy is optimal under different ecological conditions. Mounting and investing in resistance traits are accompanied by energetic expenditures of immune activation and are associated with immunopathology (immune cell damage to host tissues) (Knutie et al., 2017). These costs may manifest as increased weight loss during infection (Jog et al. 2022; Kramp et al. 2024), lower body condition (Douhard et al. 2022), and reduced growth rates (Doeschl-Wilson et al. 2009). These energetic and physiological costs can translate into reduced fitness (Miller et al. 2005), especially when not under constant threat from the pathogen, i.e., sickle cell anemia (Read et al. 2008). | By contrast, tolerant hosts endure infection without actively suppressing the parasite, which may lead to energy-expensive chronic infections (Seal et al. 2021), and some tolerance mechanisms are associated with reduced lifespan when overexpressed (Ayres and Schneider 2012). However, species known for high tolerance, such as bats, have unusually long lifespans relative to their body size, which may be attributed to their tolerance traits (e.g., wound repair) (Guth et al. 2022; Brook et al. 2023). |
| *Life History* | K-selected individuals are more likely to come into contact with a parasite to which they do not have resistance. Therefore, individuals with the ability to minimize infection-induced damage would benefit when infection is unavoidable. Species with a faster pace of life, early reproduction, large broods, and short lifespans would have less exposure to parasites throughout their lifetime. Therefore, resistance strategies, such as increased innate immune responses, could enable individuals to maintain reproductive potential without incurring costly long-term immunopathology (Sears et al., 2015). | Life-history strategies fundamentally shape how hosts invest in defense, shifting the costs and benefits of both resistance and tolerance. Species with a slower pace of life, longer lifespans, and fewer offspring may rely more on tolerance or a mixture of both strategies (i.e., adaptive immune in longer-lived animals). |
| *Environment* | It is increasingly clear that environmental factors such as temperature, resource abundance, and moisture levels profoundly affect host-parasite interactions. For example, experimental warming can cause seasonal shifts in parasite release, thereby altering when a host comes into contact with parasites. Hosts appear to exhibit seasonal variation in their resistance defenses to parasites (Rosa et al. 2022; Møller et al. 2003), which is linked to resource availability and host nutritional status (Schmid-Hempel 2003; Cotter et al. 2011; Ezenwa 2004). | Tolerance has also been shown to be labile under environmental conditions, decreasing during years of drought in Galápagos mockingbirds (McNew et al., 2019). Therefore, depending on how environmental variation affects transmissibility and parasite virulence, a disruption of annual cycles could render both tolerance and resistance less effective. |

**Concept A Supplement:**

Links to the searches for the Venn diagram containing all keywords on Web of Science.

1) Parasite load+Host defense

Key words: "parasite load" or "pathogen load" or "parasite abundance" or "parasite infection" or "parasite prevalence" or "pathogen abundance" or "pathogen prevalence" or "bd load" or "infection severity" or "increased infection" or "decreased infection" and "Host defense" or "parasite resistance" or "parasite tolerance" or "host parasite defense" or "host defence" or "host tolerance" or "host resistance"

<https://www.webofscience.com/wos/woscc/summary/9d3701da-1fa6-4cd3-a23a-cbf3522393b2-01b495ddcb/relevance/1>

2) Parasite load+ Temperature Variability

Key words: Load is the same above and "temperature variability" or "fluctuating temperature" or "variable temperature" or "thermal variability" or "thermal fluctuation" or "thermal variation" or "temperature variation"

<https://www.webofscience.com/wos/woscc/summary/aee32ed6-bdff-469a-ad97-2a641079201d-01b495f002/relevance/1>

3) Host Defense + Temperature Variability

Keywords: Same as above.

<https://www.webofscience.com/wos/woscc/summary/57d0aef9-3b82-4455-a704-d97493ee78e6-01ba0ca0bf/relevance/1>

4) Host immune + Temperature variability

Key words: "Host immune" or "innate" or "adaptive" or "immunity" or "immune" and Temperature variability if the same as above

5) Host immune + Temperature variability + Parasite load

Keywords: Combinations of the above

<https://www.webofscience.com/wos/woscc/summary/07bf8612-f7ec-4bd7-86f0-fc6b629ce2b2-01ba0c599d/relevance/1>

6) Parasite load + Host immune

<https://www.webofscience.com/wos/woscc/summary/1b11c144-467f-40a4-9b15-aca4e96977c8-01ba0c8050/relevance/1>
